# A binocular synaptic network supports interocular response alignment in visual cortical neurons

**DOI:** 10.1101/2021.06.22.449272

**Authors:** Benjamin Scholl, Clara Tepohl, Melissa A. Ryan, Connon I. Thomas, Naomi Kamasawa, David Fitzpatrick

## Abstract

In the visual system, signals from the two eyes are combined to form a coherent representation through the convergence of synaptic input populations onto individual cortical neurons. As individual synapses originate from either monocular (representing one eye) or binocular (representing both eyes) cortical networks, it has been unclear how these inputs are integrated coherently. Here, we imaged dendritic spines on layer 2/3 binocular cells in ferret visual cortex with *in vivo* two-photon microscopy to examine how monocular and binocular synaptic networks contribute to the interocular alignment of orientation tuning. We found that binocular synapses varied in degree of ‘congruency’, namely response correlation between left and right eye visual stimulation. Binocular congruent inputs were functionally distinct from binocular noncongruent and monocular inputs, exhibiting greater tuning selectivity and connection specificity. Using correlative light and electron microscopy, we found no difference in ultrastructural anatomy and instead, observed strength in numbers using a simple model simulating aggregate synaptic input. This model demonstrated a predominate contribution of binocular congruent inputs in sculpting somatic orientation preference and interocular response alignment. Our study suggests that, in layer 2/3 cortical neurons, a binocular network is responsible for forming a coherent representation in individual neurons through recurrent intracortical interactions.

## Introduction

Neural circuits in the central nervous system combine independent sources of sensory information to form coherent percepts and guide motor actions. In the visual system, a critical step in building coherent neural representations is combining signals from the two eyes. Coherent binocular convergence is the basis of cyclopean perception, stereopsis, and increased acuity and sensitivity (Barendregt et al., 2015; Parker, 2007; Scholl et al., 2013a). In carnivores and primates, binocular convergence first occurs in the primary visual cortex (V1) where individual cortical neurons respond selectively to sensory stimulation of one or both eyes (Hubel and Wiesel, 1962, 1965; Ohzawa and Freeman, 1986a; Priebe, 2008). Cortical neurons are also selective for edge orientation (Hubel and Wiesel, 1962; Priebe and Ferster, 2012), and in all mammals investigated, most binocular neurons exhibit matched orientation preferences for stimuli viewed through each eye (Bridge and Cumming, 2001; Chang et al., 2020; Ferster, 1981; Hubel and Wiesel, 1962; Nelson et al., 1977; Skottun and Freeman, 1984; Wang et al., 2010). While interocular alignment of response properties is considered to be a prerequisite for binocular vision (Marr and Poggio, 1979), the synaptic basis of this phenomenon is poorly understood.

Ultimately, the interocular alignment evident in the responses of individual neurons reflects the population of excitatory inputs synapsing onto their dendritic arbors. Receptive field properties resulting from the activity of synapses driven by the contralateral eye must be matched with those driven by the ipsilateral eye. In the simplest case, this can be conceptualized as the integration of separate, matched monocular streams converging on the same postsynaptic neuron. Experimental studies and theoretical models of interocular response alignment typically focus on this monocular perspective, proposing that contra and ipsi eye driven synapses with strong, correlated sensory-driven activity are strengthened and/or maintained during development (Bhaumik and Shah, 2014; Bienenstock et al., 1982; Erwin and Miller, 1998; Gu and Cang, 2016; Hofer et al., 2009; Mrsic-Flogel et al., 2007; Sarnaik et al., 2014; Smith and Trachtenberg, 2007; Tan et al., 2020; Wang et al., 2010; Xu et al., 2020).

Overlooked from this monocular framework, however, is the fact that much of the synaptic input a cortical neuron receives during stimulation of either eye arises from other cortical neurons that are binocular, especially in superficial V1 (Anzai et al., 1999; Ohzawa and Freeman, 1986b; Scholl et al., 2013b; Skottun and Freeman, 1984). Thus, for most binocular cortical neurons, synaptic inputs derived from both monocular and binocular neurons could play a role in interocular response alignment. While whole-cell electrophysiology is traditionally used to measure monocular and binocular excitatory synaptic conductances (Gu and Cang, 2016; Longordo et al., 2013; Scholl et al., 2013b; Zhao et al., 2013), this technique cannot disentangle the response properties of individual synaptic inputs. Consequently, the specific contributions of monocular and binocular inputs to interocular response alignment remains unresolved.

To address this issue, we used *in vivo* two-photon microscopy to visualize layer 2/3 neurons in ferret V1 expressing GCaMP6s (Scholl et al.; Wilson et al., 2016). We characterized the ocular class and orientation preference of individual dendritic spines, the postsynaptic site of excitatory synapses on pyramidal neurons. We measured responses to stimulation of either eye and compared this to the somatic output. Individual neurons with binocularly aligned somatic responses received both monocular and binocular inputs. Surprisingly, binocular inputs exhibited a varying degree of interocular alignment, quantified by the cross-correlation between each eye’s orientation tuning. We refer to this interocular correlation as ‘congruency’. Binocular congruent inputs were functionally distinct from binocular noncongruent inputs as well as monocular inputs, exhibiting greater orientation selectivity and preferences that were well matched to somatic output. A simple simulation of synaptic integration incorporating stimulus-driven activity demonstrated a predominate role for binocular congruent inputs in shaping the interocular response alignment of cortical neurons. Finally, we found that somatic interocular response alignment, across our entire population of cells, was predicted by properties of their binocular synaptic network, but not monocular synaptic networks. Our results emphasize a critical role for binocular network interactions in shaping the interocular response alignment of cortical neurons.

## Results

### Individual layer 2/3 cortical neurons receive synaptic inputs with diverse ocular properties

To investigate how monocular and binocular synaptic networks contribute to ocular alignment, we performed *in vivo* two-photon calcium imaging of layer 2/3 cells in ferret visual cortex sparsely labeled with GCaMP6s (Fig. 1a, see methods). Ferrets were visually-mature with at least 1 week of visual experience. For each cell, we optimized visual stimuli while imaging the soma. We then serially imaged visible dendritic spines on apical and basal dendrites during visual stimulation. For each imaging location, we used drifting gratings of different orientations presented to each eye independently and analyzed their responses to characterize orientation tuning (Fig. 1 b; see Methods). In total we characterized 5923 visually responsive (see Methods) dendritic spines from 35 cells.

**Figure 1:**
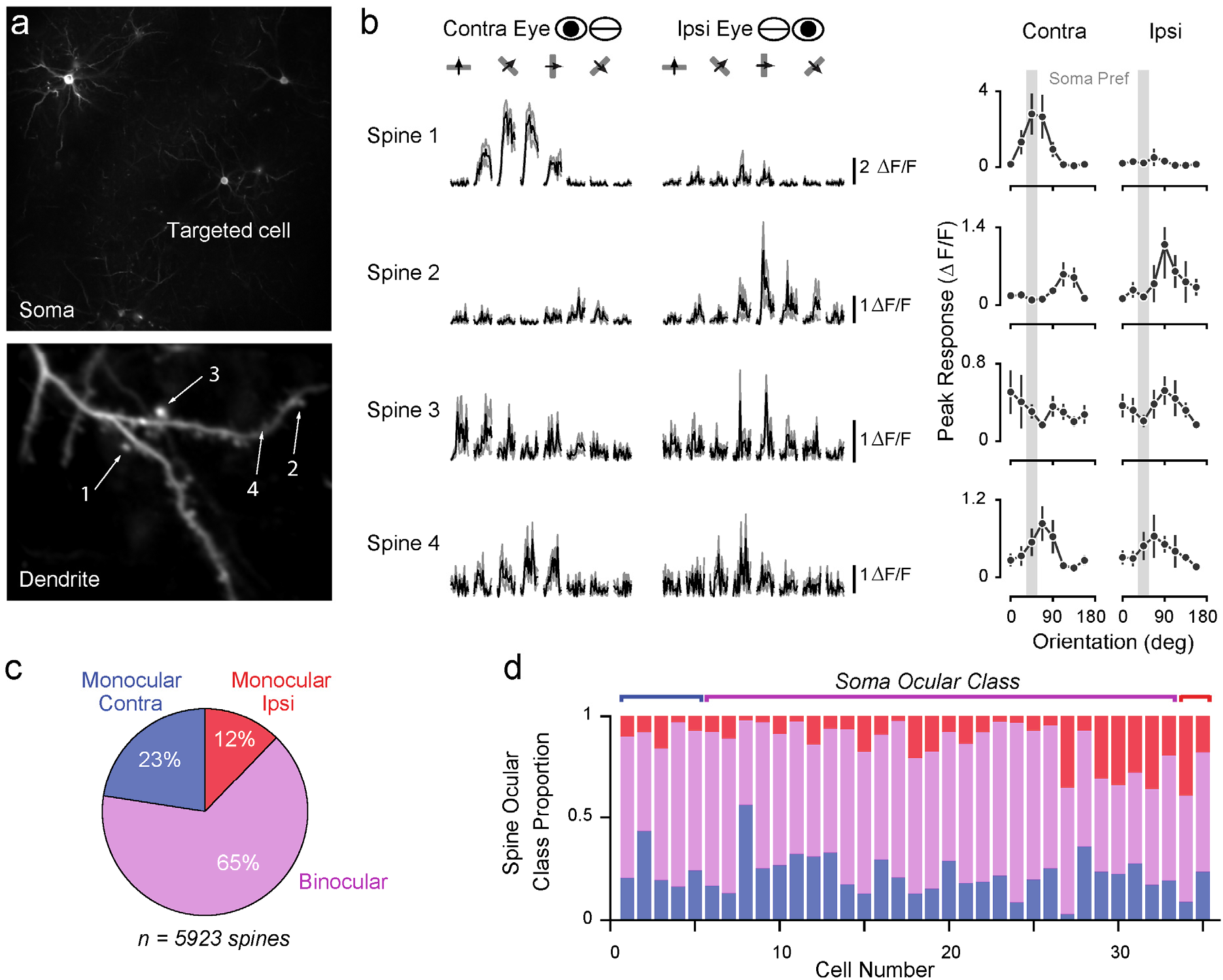
Mapping synaptic input ocularity in layer 2/3 cortical neurons with two-photon imaging of dendritic spines. **(a)** Sparse labeling of neurons in ferret V1 with GCaMP6s. Standard deviation projection of an example cell (top) and one of its dendrites (bottom) with visible spines. **(b)** Responses of four example spines shown in (a) to drifting gratings presented to the contra and ipsi eye. Shown are mean (black) and standard deviation (gray) ΔF/F responses to each stimulus (n = 8 trials) presented to the contra (left) and ipsi (right) eyes. Also shown (right column), are the peak amplitude response mean (black dots) and standard deviation (lines). The preferred orientation of the soma (soma pref) is indicated as a vertical gray bar. **(c)** Proportions of spines of each ocular class. **(d)** Proportions of spines from each ocular class for individual neurons. Same color scheme as in (c).

Ocular classes for each soma and dendritic spine were based on whether a significant response (see Methods) was elicited by visual stimulation of only the contralateral eye (contra), only the ipsilateral eye (ipsi), or both eyes (binocular). A majority of dendritic spines were classified as binocular (Fig. 1c) and a similar trend was found for individual cells (Fig. 1d, mean = 64 ± 11% s.d.). All cells examined received synaptic inputs from each ocular class, displaying a functional diversity of synaptic populations that has been reported for other visual features, sensory areas, and species (Scholl and Fitzpatrick, 2020).

In this study, we aim to determine how synaptic populations with distinct ocular properties contribute to aligned responses at the soma. Thus, we focus exclusively on binocular cells, as defined by somatic responses (n = 28/35).

### Binocular synapses vary in degree of congruency

Individual binocular synapses varied in the degree of similarity in orientation tuning between contra and ipsi eye stimulation. To characterize contra-ipsi similarity, we computed the Pearson correlation coefficient between responses evoked by stimulation each eye. Henceforth we define this metric as the degree of ‘congruency’. Two binocular example spines are shown in Fig. 1b for comparison (spine 3 and spine 4, congruency = 0.16 and 0.84, respectively). Binocular inputs exhibited a wide range of congruency, but were biased towards positive values (median = 0.38, IQR = 0.67, n = 3875; Fig. 2a). For subsequent comparisons, we split these synapses into two synaptic groups: binocular congruent (*r_C-I_* > 0.5, n = 1573 spines, 41% of all binocular inputs) and binocular noncongruent (*r_C-I_* < 0.5, n = 2302 spines, 59% of all binocular inputs). Splitting binocular synapses into two groups will be important for later analyses, as we will show that binocular congruent inputs have distinct functional properties and play a unique role in determining a cell’s feature alignment. We note that the exact cutoff value does not affect our results and, for transparency, also report analyses without this categorization.

**Figure 2:**
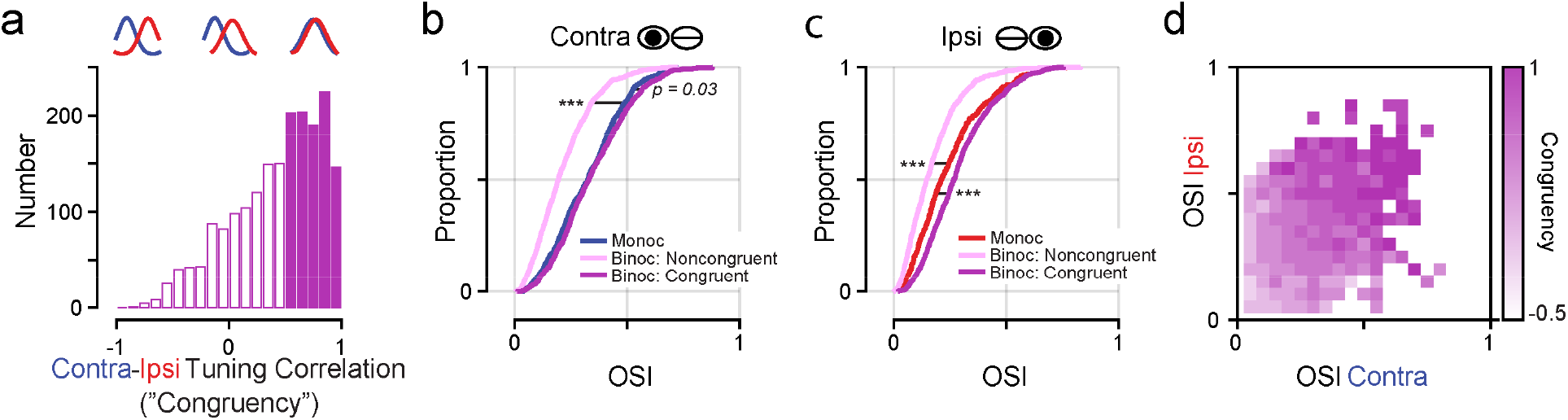
Binocular congruent inputs are most selective for orientation. **(a)** Distribution of binocular input ‘congruency’: correlation coefficient between contra and ipsi orientation tuning. Binocular inputs with a correlation >0.5 are defined as congruent (filled bars). **(b-c)** Cumulative distribution of orientation selectivity index (OSI) for monocular, binocular noncongruent, and binocular congruent synapses for contra (b) and ipsi (c) stimulation. Note, monocular contra inputs are more selective than monocular ipsi inputs (p = 0.0001, Wilcoxon ranksum test). **(d)** Heatmap of mean congruency of binocular inputs with respect to contra and ipsi OSI.

Our goal is to examine how monocular and binocular synaptic networks contribute to interocular response alignment in binocular cells. However, our dataset includes binocular cells with poor alignment, similar to previous reports in visually-mature animals (Bridge and Cumming, 2001; Chang et al., 2020). Thus, we focus exclusively on binocular cells with congruent (*r_C-I_* > 0.5) somatic orientation tuning between the two eyes (n = 16/28) and their synapses (n = 2933, binocular spines n = 1920, median congruency = 0.5) for the remainder of this study. Overall, congruency was directly related to interocular orientation preference difference, a measure used in previous studies of visual cortical neurons (circular-linear correlation = 0.74, p < 0.0001, n = 2558 binocular spines with selective tuning for each eye (see Methods)). Based on this relationship, our chosen cutoff (*r_C-I_* = 0.5) equates to ~19 deg of orientation preference mismatch.

### Binocular congruent synapses are highly orientation selective and functionally specific

We first investigated the response characteristics of the synapses for each ocular class (monocular, binocular congruent and binocular noncongruent), focusing on response selectivity and connection specificity. These two properties provide a measurement of how well a synapse conveys a given stimulus feature (selectivity) and how well synapses align to the orientation preference of the somatic output (specificity). Both properties are key factors in the interocular alignment of synaptic populations within a neuron. Additionally, we consider these properties for stimulation of the contra- and ipsilateral eye separately for the three ocular types. This allows a straightforward comparison of the properties of monocular and binocular inputs driven by the contra- and ipsilateral eyes.

Binocular congruent inputs exhibited the greatest degree of orientation selectivity (see Methods) for responses elicited by contra (Fig. 2b, median OSI = 0.33) and ipsi stimulation (Fig. 2b, c, median OSI = 0.27). This ocular class was more selective than binocular noncongruent inputs (contra: median OSI = 0.19, p < 0.0001, Wilcoxon ranksum test; ipsi: median OSI = 0.15, p < 0.0001, Wilcoxon ranksum test), monocular contra (median OSI = 0.30, p < 0.0001, Wilcoxon ranksum test), and monocular ipsi inputs (median OSI = 0.19, p < 0.0001, Wilcoxon ranksum test). Notably, binocular noncongruent inputs were the least selective for orientation overall and monocular contra inputs were more selective than monocular ipsi inputs (p < 0.0001, Wilcoxon ranksum test). There was also a strong positive correlation between congruency and selectivity for contra and ipsi stimulation (contra: Spearman’s r = 0.47, p<0.0001; ipsi: Spearman’s r = 0.50, p<0.0001; Fig. 2d).

To evaluate connection specificity, we measured the mismatch between the preferred orientation of each spine to that of their corresponding somatic output (Fig. 3a); a mismatch closer to 0 equates to greater specificity. For this analysis we excluded spines unselective for orientation (OSI < 0.10). Binocular congruent inputs were most aligned to the somatic output (contra circular median = 10.6 deg, ipsi circular median = 11.5; Figs. 3b-c). These inputs exhibited less mismatch than monocular ipsi inputs (circular median = 22.5 deg, p < 0.0001, circular Kruskal-Wallis test). A similar, but nonsignificant trend, was found for monocular contra inputs (circular median = 12.8 deg, p = 0.18, circular Kruskal-Wallis test). Binocular noncongruent inputs, on the other hand, exhibited the most mismatch for contra stimulation (circular median = 18.0 deg), significantly less than corresponding monocular inputs (p < 0.0001, circular Kruskal-Wallis test). For ipsi stimulation, binocular noncongruent inputs were indistinguishable from monocular inputs (circular mean = 20.8 deg, p = 0.62, circular Kruskal-Wallis test). Consistent with the differences between binocular categories, mismatch was inversely correlated with congruency (contra: circular-linear r = −0.30, p < 0.0001; ipsi: circular-linear r = −0.34, p < 0.0001). Remarkably, monocular contra inputs displayed less mismatch than monocular ipsi inputs (p = 0.001, Circular Kruskal-Wallis test; Fig. 3b-c), highlighting a difference between monocular input streams. In sum, we find that binocular congruent synapses are most selective and functionally specific, suggesting they are best positioned to support the binocularly aligned responses of layer 2/3 neurons.

**Figure 3:**
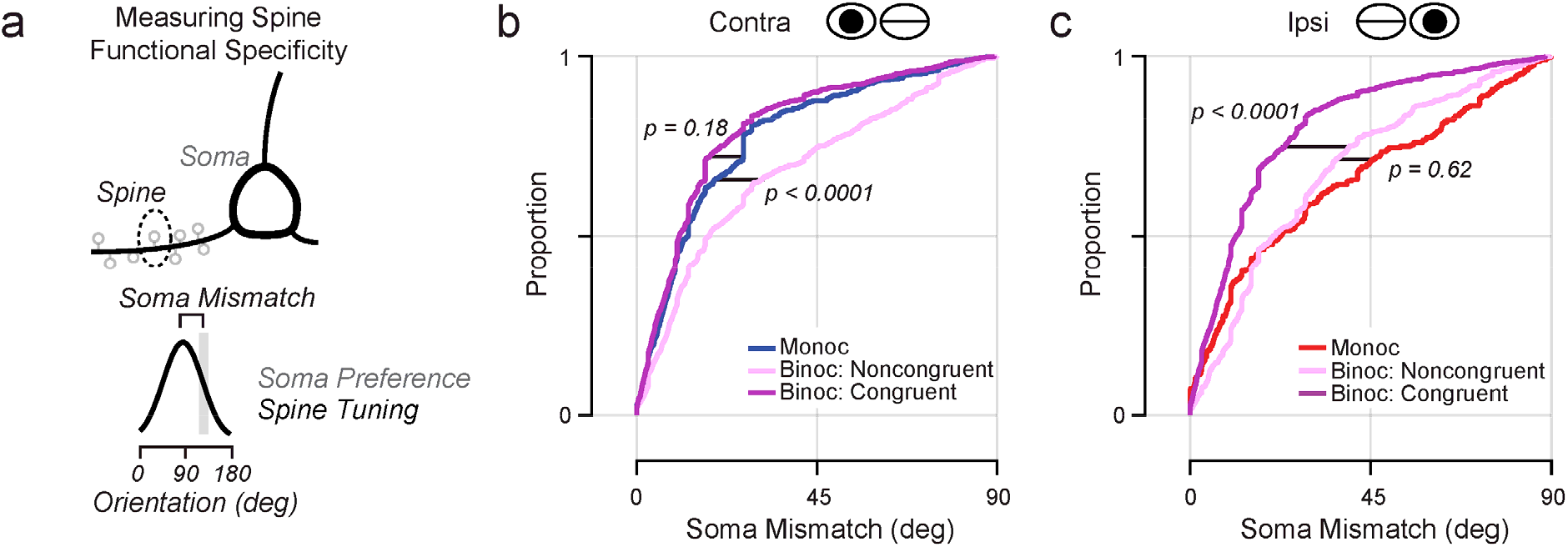
Binocular congruent inputs exhibit highest degree of functional specificity. **(a)** Functional specificity measured by comparing mismatch in orientation preference between individual spines and the soma. **(b-c)** Cumulative distribution of soma mismatch for monocular, binocular noncongruent, and binocular congruent inputs for contra (b) and ipsi (c) stimulation.

### Monocular and binocular synapses are similar in ultrastructural anatomy

While the distinct functional properties of binocular congruent inputs could support interocular response alignment, they comprise only ~1/3 of cell’s total inputs (binocular congruent: 33 ± 18 %, binocular noncongruent: 32 ± 11 %, monocular: 35 ± 13 %; proportion mean ± s.d.). However, the impact of binocular congruent synapses could be enhanced by increased strength. As synaptic strength is reflected in the ultrastructural anatomy (Holler et al. 2021; Lee et al., 2016; Scholl et al. 2021), one possibility is that binocular congruent synapses also share distinct ultrastructural properties.

We recently developed a method to correlate light and electron microscopy at the synaptic level (Fig. 4; see Methods), allowing comparison of the same synapses (Thomas et al., 2020). In a subset of the binocular congruent cells (n = 4) imaged *in vivo* and examined in this study, we anatomically reconstructed dendritic spines from basal dendrites. In a subset of these spines (n = 103) we classified ocularity based on the criteria used in this study (see Methods). From this dataset, we examined 3 basic anatomical features: spine head volume, postsynaptic density (PSD) area, and spine neck length (Fig. 4d). Spine head volume and PSD area are positively correlated with synapse strength (Bourne and Harris, 2011; Toni et al., 1999), while spine neck length can attenuate synapse strength (Araya et al., 2006).

**Figure 4:**
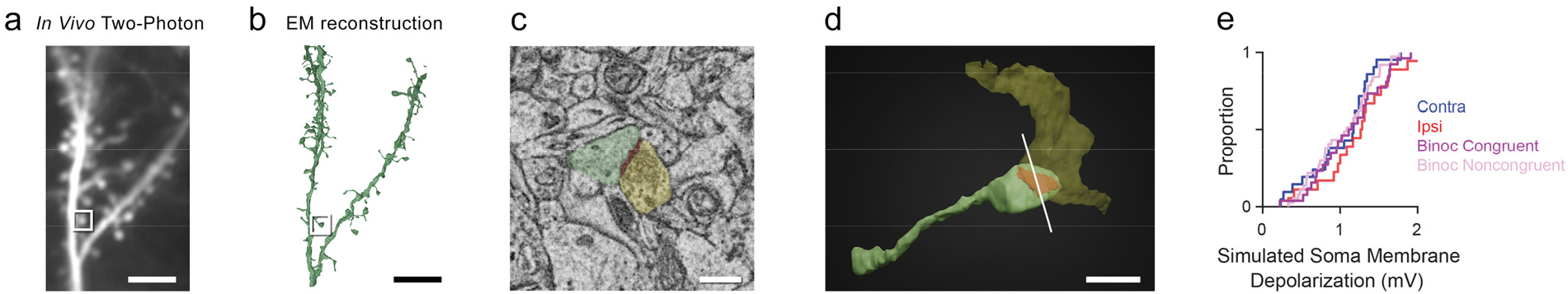
Correlation of light and electron microscopy at synaptic resolution. **(a)** Example in vivo two-photon standard deviation projection of dendrite and spines. Scale bar is 10 μm. **(b)** Same dendrite and spines reconstructed from an electron microscopy volume. Scale bar is 10 μm. **(c)** Electron micrograph of an example synapse in (a-b) showing the presynaptic bouton (yellow), postsynaptic density (red), and dendritic spine head (green). Scale bar is 500 nm. **(d)** Full 3D reconstruction of the example synapse shown in (a-c). The white bar represents the cross section in (c). Scale bar is 1 μm. **(e)** Cumulative distributions of simulated soma membrane potential depolarization for synapses of each ocular class. Postsynaptic anatomical measurements were combined in a NEURON model to simulate effective somatic input (see Methods).

Overall, binocular congruent inputs were similar in ultrastructure compared to other ocular classes (Table 1; Supp. Fig. 1). This was supported by a nonparametric one-way analysis of variance of spine head volume (p = 0.58), spine PSD area (p = 0.098), and spine neck length (p = 0.093). We also found no relationship between congruency and any other anatomical feature (spine head volume: Spearman’s r = 0.003, p = 0.49; PSD area: Spearman’s r = 0.10, p = 0.21; neck length: Spearman’s r = −0.009, p = 0.47). We found similar results when considering a metric of synaptic strength that combines multiple anatomical features and their interactions (e.g. synapse PSD area and neck length inversely regulate synapse strength). We used a NEURON model of each synapse, incorporating all anatomical features, and simulated the somatic depolarization due to a single action potential (see Methods). This showed that binocular congruent inputs were similar in effective strength as other ocular classes (p = 0.50, nonparametric one-way ANOVA; Table 1; Fig. 4e). Likewise, congruency was unrelated to simulated somatic input amplitude (Spearman’s r = 0.07, p = 0.71). Altogether, our data demonstrate that binocular congruent synapses are similar in ultrastructural anatomy and simulated effective strength compared to monocular and noncongruent synapses.

**Table 1:** Ultrastructural properties of synapses of each ocular class. Values reported are mean and standard deviation for each ocular class and anatomical feature or metric of strength.

| **Class** | **Spine Head Volume ($\mu\text{m}^3$)** | **Spine PSD Area ($\mu\text{m}^2$)** | **Spine Neck Length ($\mu\text{m}$)** | **Simulated Soma Input (mV)** |
| --- | --- | --- | --- | --- |
| Monocular Contra<br>(*n* = 21) | 0.35 ± 0.26 | 0.26 ± 0.17 | 1.97 ± 1.04 | 1.02 ± 0.42 |
| Monocular Ipsi<br>(*n* = 18) | 0.48 ± 0.29 | 0.38 ± 0.29 | 2.32 ± 1.05 | 1.25 ± 0.52 |
| Binocular congruent<br>(*n* = 26) | 0.42 ± 0.31 | 0.37 ± 0.25 | 1.78 ± 0.82 | 1.12 ± 0.44 |
| Binocular noncongruent<br>(*n* = 38) | 0.40 ± 0.32 | 0.24 ± 0.19 | 1.72 ± 0.62 | 1.04 ± 0.40 |

### Simulating synaptic population activity and the emergence of interocular aligned responses

Binocular congruent synapses have distinct response properties suggesting they are uniquely positioned to support interocular alignment, but their ultrastructural features (a measurement of strength) are largely similar to other ocular classes. Strength alone, however, cannot predict how synapses might drive somatic activity during visual stimulation. Somatic activity derives from the net pattern of activity of functionally diverse populations of monocular and binocular synapses that are recruited by a particular stimulus in a stochastic manner.

To understand how activity patterns of monocular and binocular synapses shape alignment, we devised a simple simulation of the aggregate synaptic drive onto each neuron (drawn from each neuron’s measured synaptic population). Our simulation incorporates only synaptic calcium activity (converting ΔF/F into binary events) during presentations of visual stimuli (example branch shown in Figure 5a). For each simulation iteration, a random trial of one set of stimuli was chosen and the activity across all dendrites and all groups of spines was summed to create an aggregate, simulating somatic aggregate input for each cell (Fig. 5b-d). This procedure was repeated 10,000 times for contra and ipsi visual stimulation. Simulation of the total aggregate for a representative binocular congruent cell is shown in Figure 5d (n = 197 spines). While we ignored factors such as dendritic compartmentalization and active nonlinearities, this simulation provides an estimate of stimulus-driven input to the soma from each ocular class, incorporating sensory-driven activity, synapse stimulus selectivity, functional specificity, and reliability of activation across trials.

**Figure 5:**
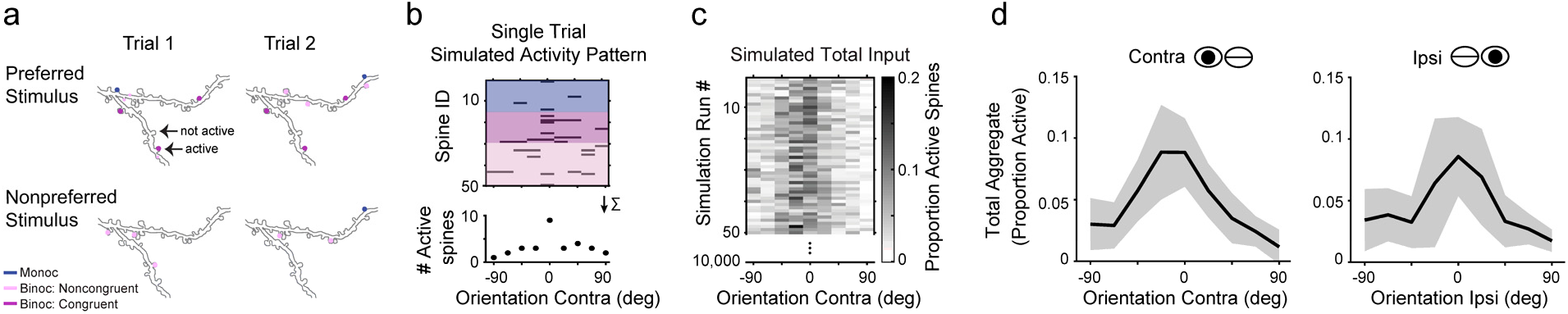
A simple model simulating synaptic input aggregates. **(a)** Example trial-to-trial activity of dendritic spines at the somatic preferred and nonpreferred stimulus. Dendritic spines filled with color are those active. **(b)** Activity pattern (discrete calcium events, shown in black, see Methods) of a subset of spines (n = 50) for a single simulation iteration. Shown is activity driven by contra stimulation, with respect to the somatic preferred orientation. Spines are grouped and color-coded by their ocular class. To estimate total aggregate input for a single trial, the activity of all spines is summed (*bottom*). **(c)** Individual total aggregates for 50 (out of 10,000) simulation runs. Data is the proportion of possible spines active (here 197 spines). **(d)** Total synaptic aggregate estimated for contra and ipsi stimulation. Shown is the mean and standard deviation.

Across binocular congruent cells (n = 14, total spines = 192 ± 69, mean ± s.d.), the proportion of active spines was greatest at the somatic preferred orientation (contra: 18.6 ± 2.2%, ipsi: 11.5 ± 2.1%, mean ± s.e.; Fig. 6a). At the somatic preferred orientation, the proportion of active synapses from each group was similar (binocular congruent: 6.5% and 5.4%, binocular noncongruent: 5.9% and 4.8%, monocular: 6.1% and 3.9%, mean proportion active for contra and ipsi eyes; Fig. 6a). In contrast, for nonpreferred orientations, monocular and binocular noncongruent inputs were recruited more readily (binocular congruent: 1.4% and 1.3%, binocular noncongruent: 3.0% and 2.8%, monocular: 2.5% and 2.6%, mean proportion active for contra and ipsi eyes; Fig. 6a).

**Figure 6:**
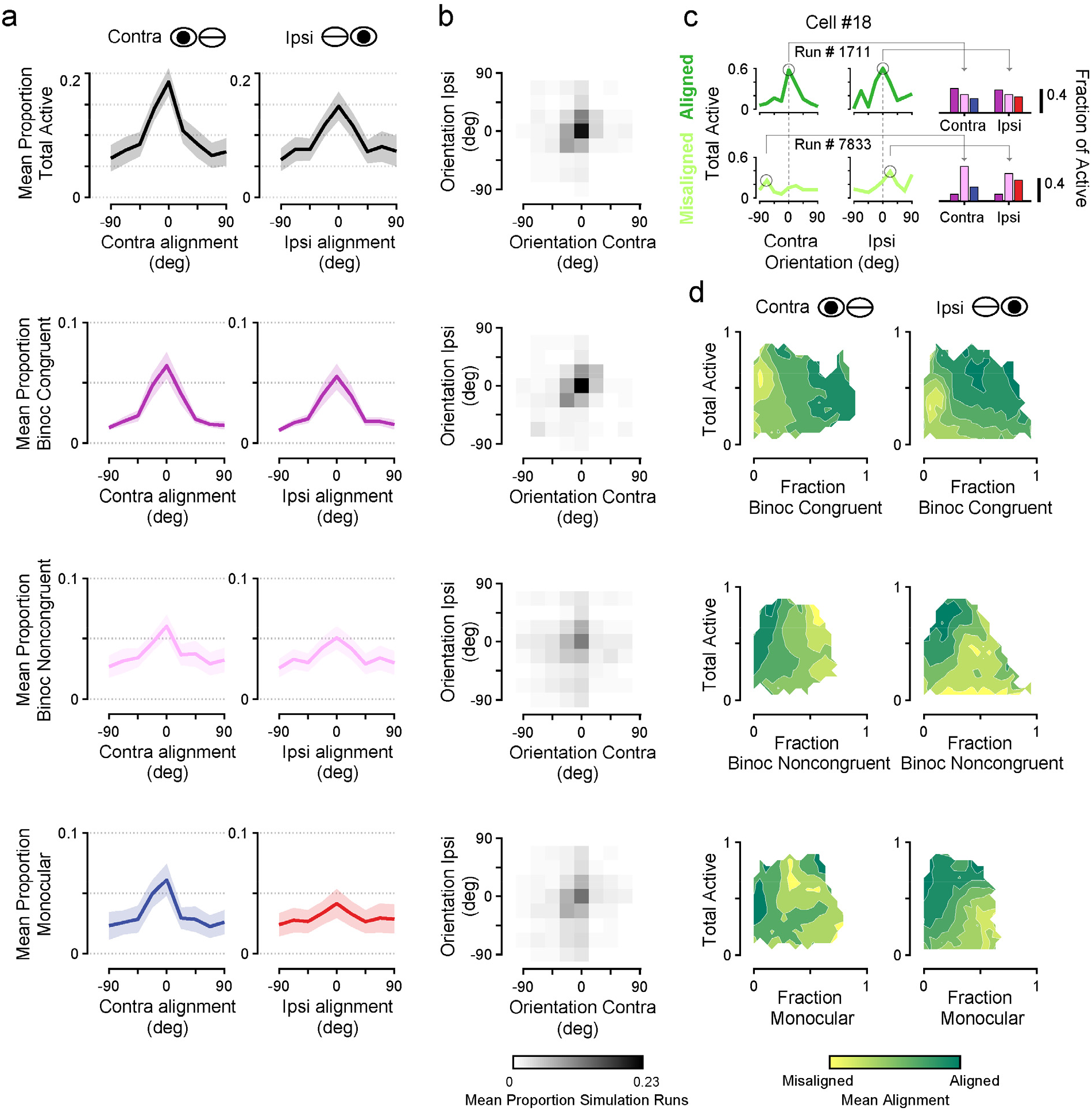
Interocular alignment of total synaptic aggregates strongly depend on activation of binocular congruent synapses. **(a)** Simulated aggregate tuning for contra and ipsi stimulation across 14 binocular congruent cells. From top to bottom: total aggregate, binocular congruent populations, binocular noncongruent populations, and monocular populations. Tuning is with respect to the soma preferred orientation; data are mean and standard error. **(b)** Interocular distributions of trial-to-trial alignment of aggregates to the somatic preferred orientation in contra and ipsi stimulation conditions. Each bin is frequency across simulation iterations. **(c)** Measurement of active synapses with respect to the total aggregate interocular alignment. *Left:* Shown is the proportion of spines active for each orientation (with respect to somatic orientation preference) and eye condition for two example simulation runs: one aligned (dark green) and one misaligned (light green). Circles denote maximum number of active synapses and orientations used to calculate alignment. *Right:* the fractions of spines from each ocular class contributing to the maximum values. Same color scheme as in (a). **(d)** Mean alignment conditioned on the proportion of total synapses active (ordinate, normalized by the maximum number active observed) and fraction of those from each ocular class (abscissa). Data are mean across all simulation runs from all congruent cells.

We next assessed the degree of alignment in each ocular population by examining distributions of interocular orientation preference (relative to somatic preferred) across simulation iterations. Preferred orientation was defined as the stimulus evoking the largest proportion of active spines, similar to a maximum a posteriori estimate (Fig. 6b). Binocular congruent aggregates were more frequently aligned than monocular or noncongruent aggregates (p = 0.006 and p = 0.018, respectively, sign-rank one-sided test; Fig 6b). While frequency of alignment for all ocular classes was correlated with the total aggregate (monocular: Spearman r = 0.75, p < 0.0001, binocular congruent: Spearman r = 0.93, p < 0.0001, binocular noncongruent: Spearman r = 0.82, p < 0.0002, one-sided tests), binocular noncongruent and monocular aggregates were consistently *less* aligned than the total (p = 0.002 and p = 0.01, respectively, sign-rank one-sided test). In contrast, binocular congruent aggregates were indistinguishable from the total aggregate (p = 0.33, sign-rank one-sided test, mean difference = −0.01 +/− 0.06 s.d.).

Thus far, our simulation suggests that (1) a greater number of active synapses contribute to the somatic preferred orientation for both eyes and (2) the binocular congruent synapse population exhibits a degree of alignment most similar to the total population. These analyses, however, do not consider variation across each simulation run, nor the relationship between the number of active synapses and degree of alignment. Further, as the total aggregate (Fig. 6a, *top*) is most akin to subthreshold input at the soma, we wanted to understand how the number of active synapses from each ocular class contributes to the total aggregate and its alignment. To address this issue, we measured the total aggregate alignment (defined as the distance from the somatic preferred orientation) and the total number of active synapses at the preferred orientations (Fig. 6c, black circles) for each simulation iteration. We then examined the contribution of different ocular classes of synapses at preferred orientations (Fig. 6c, *right*). As shown for individual simulation runs in Figure 6c, an aligned total aggregate had a large number of active synapses at the somatic preferred orientation and binocular congruent synapses were the largest contributor. In contrast, a misaligned total aggregate received more input from binocular noncongruent and monocular synapses and showed less activation overall.

Across our population of cells, we found that when the total number of active synapses is large and a fraction of those are binocular congruent, the total aggregate is more aligned (Fig. 6d, *top*). For binocular noncongruent synapses, we observed a different relationship: alignment correlated with fewer active binocular noncongruent synapses and greater activation of the total population (Fig. 6d, *middle*). For monocular ipsi synapses, we observed a similar relationship as for binocular noncongruent synapses (Fig. 6d, *bottom right*), and the fraction of monocular contra synapses recruited showed little relationship to alignment or the total synapses active (Fig. 6d, *bottom left*). Multivariate linear regression models largely confirmed these observations (Table 2, see Methods), but for contra visual stimulation, the total number of active synapses was a weak predictor. These relationships were also found for individual cells; the total number of active spines was positively related to alignment in a majority of cells (contra: binocular congruent = 10/14, binocular noncongruent = 9/14, monocular = 10/14; ipsi: binocular congruent = 14/14, binocular noncongruent = 14/14, monocular = 14/14 cells). With respect to each ocular class, a majority of cells showed that the fraction of binocular congruent spines was positively related to alignment (contra: 8/14; ipsi: 9/14), while binocular noncongruent and monocular fractions were inversely related to alignment (contra: 10/14 and 9/14, respectively; ipsi: 9/14 and 12/14, respectively). Interestingly, if an interaction term was included in the linear regression model, it was weighted more heavily (Table 3), providing further evidence that alignment is predicted by the total number of active synapses and total fraction of those that are binocular congruent.

**Table 2:**
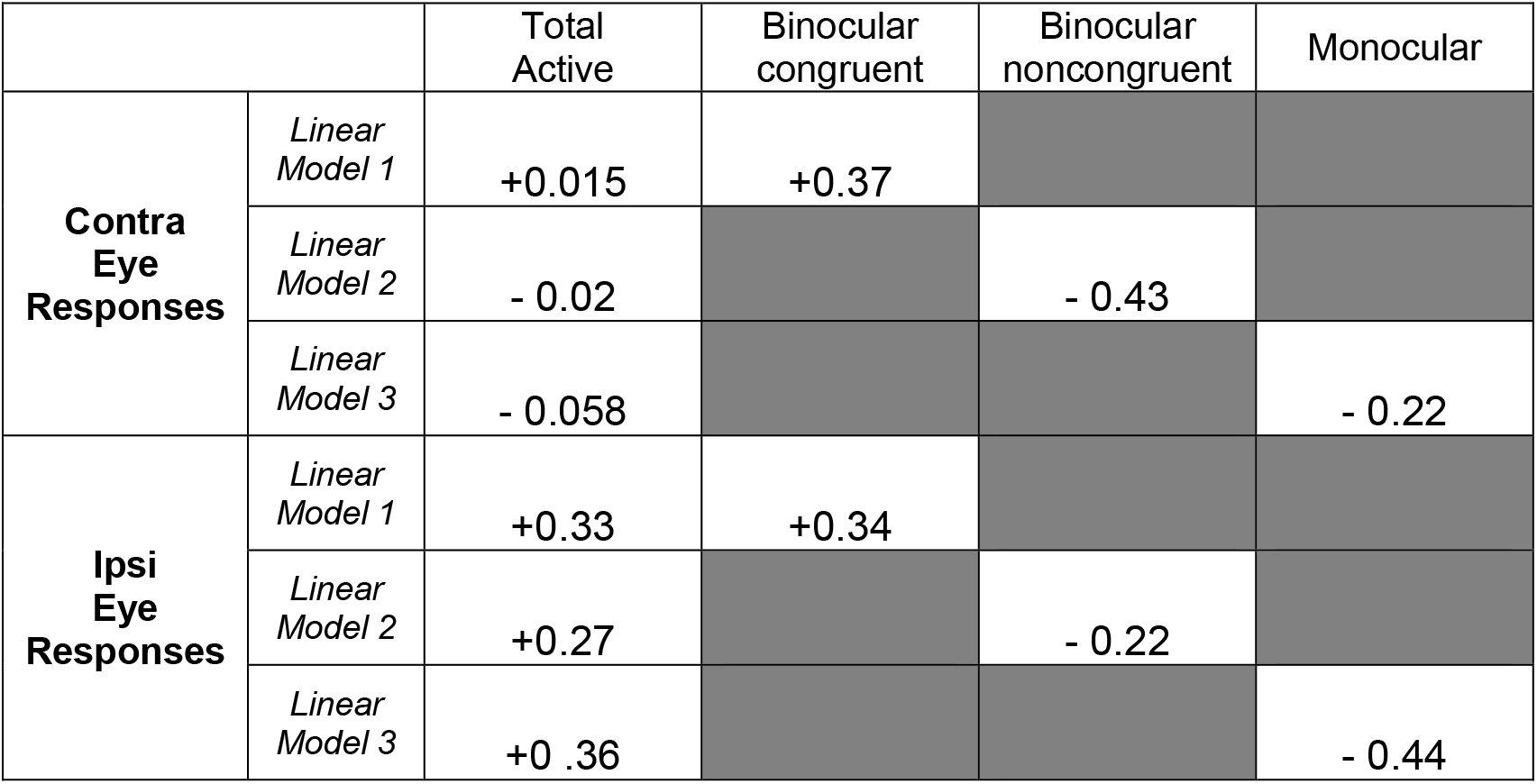
Linear regression model predicting simulated total aggregate interocular alignment. Shown are coefficients from model fits. All coefficients were statistically significant (p < 0.05). Coefficients for noise model and constant terms (see Methods) were not significant.

|  |  | Total Active | Binocular congruent | Binocular noncongruent | Monocular |
| --- | --- | --- | --- | --- | --- |
| **Contra Eye Responses** | *Linear Model 1* | +0.015 | +0.37 |  |  |
|  | *Linear Model 2* | - 0.02 |  | - 0.43 |  |
|  | *Linear Model 3* | - 0.058 |  |  | - 0.22 |
| **Ipsi Eye Responses** | *Linear Model 1* | +0.33 | +0.34 |  |  |
|  | *Linear Model 2* | +0.27 |  | - 0.22 |  |
|  | *Linear Model 3* | +0 .36 |  |  | - 0.44 |

**Table 3:** Linear regression model with interaction term predicting simulated total aggregate interocular alignment. Shown are coefficients from model fits. All coefficients were statistically significant (p < 0.05) unless stated otherwise. Coefficients for noise model and constant terms (see Methods) were not significant.

|  |  | Total | Binocular congruent | Binocular noncongruent | Monocular | Interaction |
| --- | --- | --- | --- | --- | --- | --- |
| **Contra Eye Responses** | *Linear Model 1* | - 0.079 | + 0.233 |  |  | + 0.29 |
|  | *Linear Model 2* | + 0.26 |  | - 0.036 |  | - 0.86 |
|  | *Linear Model 3* | - 0.064 |  |  | - 0.23 | 0.017 (*n.s.*) |
| **Ipsi Eye Responses** | *Linear Model 1* | + 0.10 | + 0.14 |  |  | + 0.59 |
|  | *Linear Model 2* | + 0.65 |  | + 0.10 |  | - 1.12 |
|  | *Linear Model 3* | + 0.48 |  |  | - 0.29 | - 0.47 |

Our simulations suggest that interocular alignment of synaptic aggregates depends on the contribution of binocular congruent populations and the total number of active synapses. While binocular congruent inputs do not constitute the majority of inputs to a cell overall, they are selectively recruited at the somatic preferred orientation, resulting in a disproportionate impact. This impact could be greatly enhanced by co-active monocular and binocular noncongruent inputs to increase synaptic drive overall, which is likely critical to drive somatic activity and increase the probability of generating spiking activity. Altogether, it is clear that binocular congruent synapses strongly support feature-matched responses between the two eyes.

### Somatic congruency is predicted by the relative proportion of binocular synaptic inputs

While we have focused solely on binocular congruent cells, it is possible that diversity in somatic congruency derives from the underlying synaptic inputs. We wondered whether characteristics of our synaptic populations could predict the degree of congruency of their respective soma; specifically, we wondered whether the number of available inputs from each ocular class could predict somatic congruency. Not surprisingly, noncongruent cells (n = 12) had significantly less binocular congruent synaptic inputs than congruent cells (18 ± 8% and 33 ± 18%, respectively, mean ± s.d., p = 0.01, one-sided Wilcoxon ranksum test). Therefore, we compared the proportion of spines from each ocular class to somatic congruency. We observed a tradeoff between congruent and binocular noncongruent synapses: the proportion of binocular congruent synapses was positively correlated with somatic congruency (r = 0.47, p = 0.006, one-sided Spearman correlation; Fig. 7a) and the proportion of binocular noncongruent synapses was inversely correlated with somatic congruency (r = −0.63, p < 0.0002, onesided Spearman correlation; Fig. 7b). In contrast, the proportion of monocular spines was unrelated to somatic congruency (contra: r = −0.07, p = 0.7, ipsi: r = 0.04, p = 0.8, twosided Spearman correlation; Fig. 7c), highlighting the unique influence of binocular populations.

**Figure 7:**
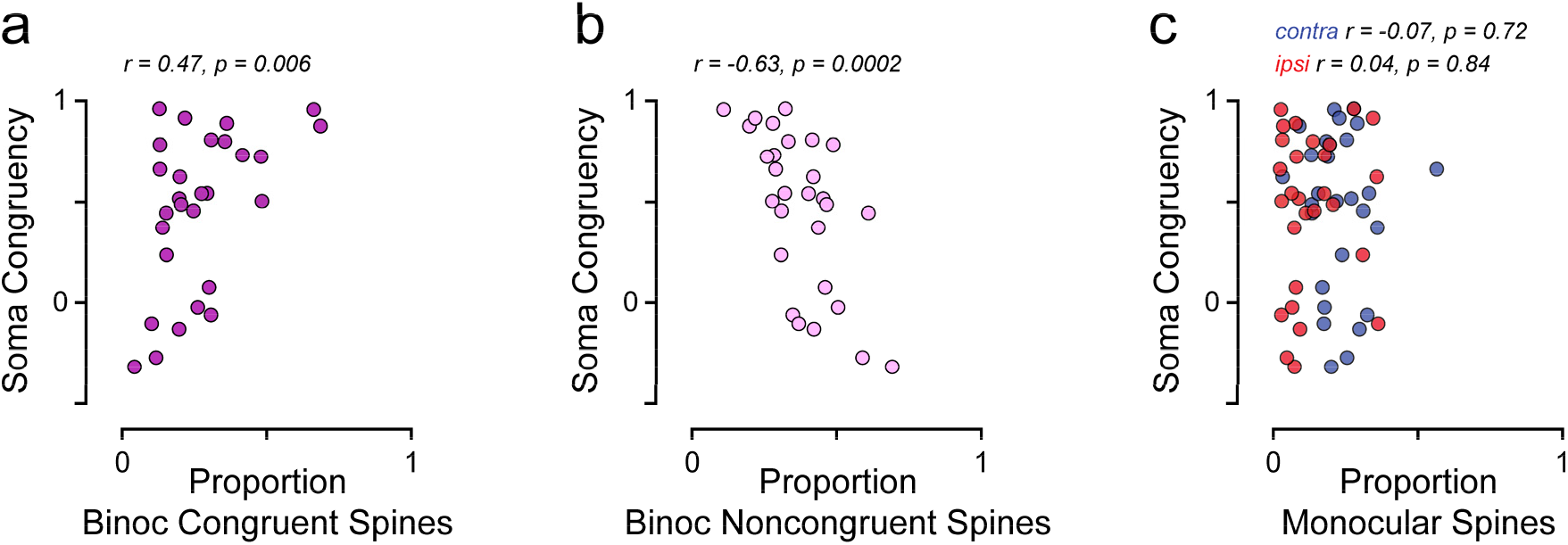
Somatic congruency correlates with synaptic input population properties. **(a)** Relationship between the proportion of binocular congruent synapses and somatic congruency. Each data point is from a single cell. (**b**) Same as in (a) for binocular noncongruent synapses. (**c**) Same as in (a) for monocular contra (blue) and monocular ipsi (red) synapses.

In sum, our results suggest that binocular congruency and interocular alignment of orientation preference lie in the balance of aligned, selective congruent and less selective, misaligned binocular noncongruent inputs.

## Discussion

Visual cortical neurons exhibit binocularly matched responses, yet it is unknown how this coherent representation emerges from converging sets of synapses which originate from both monocular and binocular neurons. In this study we examined synaptic populations on individual layer 2/3 neurons in ferret visual cortex to determine how different networks contribute to the interocular alignment of orientation preference. All cells were innervated by inputs driven by stimulation of the ipsilateral eye or contralateral eye (monocular) and inputs driven by stimulation of either eye (binocular). Binocular inputs could be split into two groups (congruent and noncongruent), depending on the degree of correlation between contra and ipsi orientation tuning. Binocular congruent inputs were distinct as they were most selective for orientation and exhibited the highest degree of connection specificity but were not different in ultrastructural anatomy compared to other input types. To better understand how monocular and binocular synaptic networks are integrated to produce an aligned response, we used a simple simulation combining sensory-driven synaptic activity. This simulation revealed a predominate role for binocular congruent inputs, contributing a selective and highly aligned aggregate input. This arose through the combination of their functional specificity, tuning selectivity, and combined strength in numbers. Finally, we observed that congruent responses at the soma of binocular cells was predicted by the relative contribution of binocular congruent and noncongruent inputs.

Our study highlights the role of a distinct binocular synaptic network in shaping interocular response alignment. As cortical binocularity emerges over development in carnivores (Casagrande and Boyd, 1996; Chang et al., 2020; Hubel and Wiesel, 1965), these networks might involve recurrent amplification, such as found for stimulus-tuned attractor networks (Ahmadian et al., 2013; Ben-Yishai et al., 1995). Visual stimuli preferred by the soma could be amplified by binocular congruent synapses, as they provide highly specific (and aligned) excitatory drive. Noncongruent inputs might provide nonspecific excitatory drive, which could act to depolarize the soma closer to spike threshold and increase the probability of eliciting action potentials (Priebe, 2008). Altogether, it is clear that at this cortical processing stage, binocular, rather than monocular, synaptic networks are most important in shaping interocular alignment of neural response properties.

### Comparison with a model of monocular convergence

Conventional models of binocularly matched responses focus on the convergence of monocular input streams. In these models, contra and ipsi inputs arriving onto single neurons become matched with one another through correlation-based plasticity mechanisms. This has been a common framework for interpreting experimental findings (Gu and Cang, 2016; Sarnaik et al., 2014; Tan et al., 2020; Wang et al., 2010). Additional support comes from models that recapitulate experimental findings by simulating interactions between monocular input streams (Bhaumik and Shah, 2014; Bienenstock et al., 1982; Erwin and Miller, 1998; Xu et al., 2020). Thus, the expectation has been that monocular input populations would exhibit a high degree of match in orientation preference. In contrast, our results from visually mature ferrets do not support this prediction, as monocular inputs did not significantly contribute to response alignment within individual layer 2/3 neurons.

It is important to point out that prior experimental studies differ from our own in a key aspect: the method of recording from synaptic populations. Previously, monocular and binocular inputs were characterized using electrophysiological measurements of subthreshold membrane potential or current with intracellular recordings (Longordo et al., 2013; Scholl et al., 2013b; Zhao et al., 2013). During contra and ipsi (monocular) visual stimulation, these measurements reflect coactivation of binocular and respective monocular synapses, so monocular and binocular synaptic networks cannot be isolated. In contrast, dendritic spine imaging captures synapses contributing to each network. Our results offer an alternative interpretation of previous experimental findings: the aligned responses to monocular stimulation are strongly driven via a binocular network, not two converging monocular networks.

Our study does not rule out a monocular convergence hypothesis entirely. Layer 2/3 neurons that we targeted primarily receive intracortical input and little or no thalamocortical innervation (Binzegger et al., 2004). It is possible that populations of thalamocortical inputs, which converge onto stellate cells in layer 4, are more in line with a model of monocular convergence. Thalamocortical synapses are almost entirely monocular (in carnivores and primates) and may have a greater influence on cortical cells as their synapses tend to be larger and have a greater number of presynaptic release sites (Ahmed et al., 1994; Kharazia and Weinberg, 1994). In addition, thalamocortical inputs could provide an initial bias in interocular alignment before requisite binocular visual experience (Chang et al., 2020; Gu and Cang, 2016) and this bias could be amplified by layer 2/3 cortical circuits, as they receive input from layer 4. Future studies will be necessary to map synaptic populations onto layer 4 neurons, and compare thalamic and cortical contributions to interocular response alignment.

### Structural properties of binocular synapses

In this study we do not find systematic differences in the ultrastructural anatomy between different ocular classes. Instead, we observe strength in numbers of synapses, similar to our previous study on orientation tuning (Scholl et al., 2020): binocular synapses were most numerous overall, followed by monocular contra- and ipsilateral inputs, and each ocular class draws from the same synaptic strength distributions. Thus, the aggregate strength of a synaptic network is largely determined by numerosity (Fig. 6). We do, however, observe some curious structure-function relationships. For example, binocular synapses, as a whole, have shorter necks than monocular synapses (Table 1). As spine neck length can regulate synaptic strength (Araya et al., 2006), this might indicate a predominate role of binocular synaptic networks in shaping layer 2/3 neuron response properties. These spines may also have more energetic demands, as binocular viewing may drive them more frequently than monocular synapses, and shorter necks place the synaptic contact closer to molecular machinery within the dendrite.

### A potential role of ‘noncongruent’ binocular inputs

Synaptic imaging continues to reveal that the networks underlying cortical neuron selectivity are more complicated and diverse than previously thought (Scholl and Fitzpatrick, 2020). In this study, we observed diversity in ocular properties. While we focused on binocular congruent inputs, we acknowledge that there is likely a role for those classified here as ‘noncongruent’. One interesting possibility is that these inputs drive neurons under binocular viewing conditions where the left and right eyes are not receiving exactly the same visual information. For example, these inputs might be responsible for signaling motion in depth (Cormack et al., 2017) or signaling features during occlusion of one eye (Anderson and Nakayama, 1994). They might also provide signals to convey a horizontal disparity (i.e. surface slant) (Blakemore et al., 1972; Greenwald and Knill, 2009) and neural models of horizontal disparity predict that the input filters would differ in orientation and be noncongruent by our measure.

Another interesting possibility is that these inputs could generate invariant binocular response properties across binocular viewing conditions. For example, head tilt can cause torsional eye movements, so visual features falling on the retina would no longer be matched along the horopter. Under these conditions, synapses we defined as ‘noncongruent’ might actually be congruent in their responses to visual features. In this way, the somatic output of the cell would be invariant to different binocular viewing conditions and could provide information under a variety of situations. Thereby, diversity may reflect the ability of visual neurons and the network as a whole to encode information invariant of viewing conditions. This could provide an explanation for our observation that binocular noncongruent inputs exhibit synaptic strengths comparable to binocular congruent inputs, as we assessed congruency in just one condition and categorized inputs for this specific situation, neglecting conditions where the ‘noncongruent’ inputs provide the strongest and most relevant input.

There is always the possibility that our limited stimulus set inaccurately characterized synapses classified as ‘noncongruent’. However, a number of studies of V1 neurons report fairly large interocular orientation preference differences (which is directly related to congruency) (Blakemore et al., 1972) from a sizeable fraction of cells. Given that an orientation preference difference of ~20 degree corresponds to our congruency cut-off (*r_c-l_* = 0.5), about 25-30% of neurons in visually-mature ferrets (Chang et al., 2020) and ~10% of neurons in adult primates (Bridge and Cumming, 2001) are classified as noncongruent. This suggests that noncongruent populations reflect more than a mere measurement error.

### Development of a binocular synaptic network supporting interocular response alignment

We propose that binocular visual experience in coordination with the structural development of cortical circuits gives rise to the binocular congruent synaptic network. Prior to visual experience or the critical period of development, cells exhibit a higher degree of interocular mismatch in orientation preference (Chang et al., 2020; Tan et al., 2020; Wang et al., 2010), but initial biases could act as starting point for later development. Initial biases could result from either (1) thalamocortical inputs (Gu and Cang, 2016), (2) spontaneous cortical activity patterns structuring receptive field properties (Smith et al., 2018), (3) random connectivity (Ko et al., 2013), or any combination of these possibilities. One study from Gu and Cang (2016) found evidence that, before the critical period in mouse, an initial bias is provided by thalamocortical input to layer 4 neurons, while intracortical input exhibits greater mismatch. After maturation, intracortical and thalamocortical input are equivalently matched, suggesting that intracortical connectivity develops, either through instruction from thalamocortical input or de novo. However, as the authors were unable to record from the same cells over time, it is not possible to distinguish between these two possibilities.

In comparing mouse and ferret studies, it is important to point out that the critical period of development in mouse begins almost 2 weeks after eye opening, and it is unknown whether this initial binocular experience shapes cortical circuits. In contrast, Chang et al. (2020) observed a dramatic shift in binocular response alignment in ferret immediately after eye opening. In addition, they found a considerable amount of congruency in binocular cells, *prior* to binocular visual experience. Thus, at least in ferret, this initial network, which could form through Hebbian modification of horizontal connections without thalamic innervation (Bartsch and van Hemmen, 2001), might instruct the maturation of a binocular congruent synaptic network.

So far, we considered only the functional maturation of cellular and synaptic networks early in development, neglecting concomitant changes in structure. Prior to and following eye opening in the developing visual cortex, dendritic arborization of pyramidal cells increases in size and complexity (Callaway and Borrell, 2011). After the onset of visual experience, long-range horizontal connections between layer 2/3 cells are established and stabilized (Durack and Katz, 1996) and synaptic density increases dramatically (Cragg, 1975; Yuste and Bonhoeffer, 2004). Increased synaptic connections lead to increased synaptic dynamics, as synaptic remodeling and elimination continues throughout development (Bhatt et al., 2009). In addition, early in development, the size (strength) of synaptic contacts increases dramatically (Cizeron et al., 2020). As structural development of cortical circuitry coincides with the onset of binocular vision, it stands to reason that binocular synaptic networks emerge and consolidate around this time. In one way, this process can be thought of as *structural recurrent amplification*; addition and stabilization of binocular intracortical inputs boosts the excitatory drive underlying interocular response selectivity and alignment. We believe this view, of a dynamic and stabilizing recurrent circuit emerging through development will provide insight into the nature of binocular synaptic networks and developmental maturation of binocular vision.

## Acknowledgements

The authors thank the Fitzpatrick lab for helpful scientific discussion, the GENIE project for access to GCaMP6s, and the Max Planck Florida Institute ARC for animal care. This work was supported by funding from NIH grants R01 EY011488 (D.F.), NIH grant K99 EY031137 (B.S.), the Max Planck Florida Institute for Neuroscience, and the Max Planck Society.

---

**Supplemental Figure 1:**
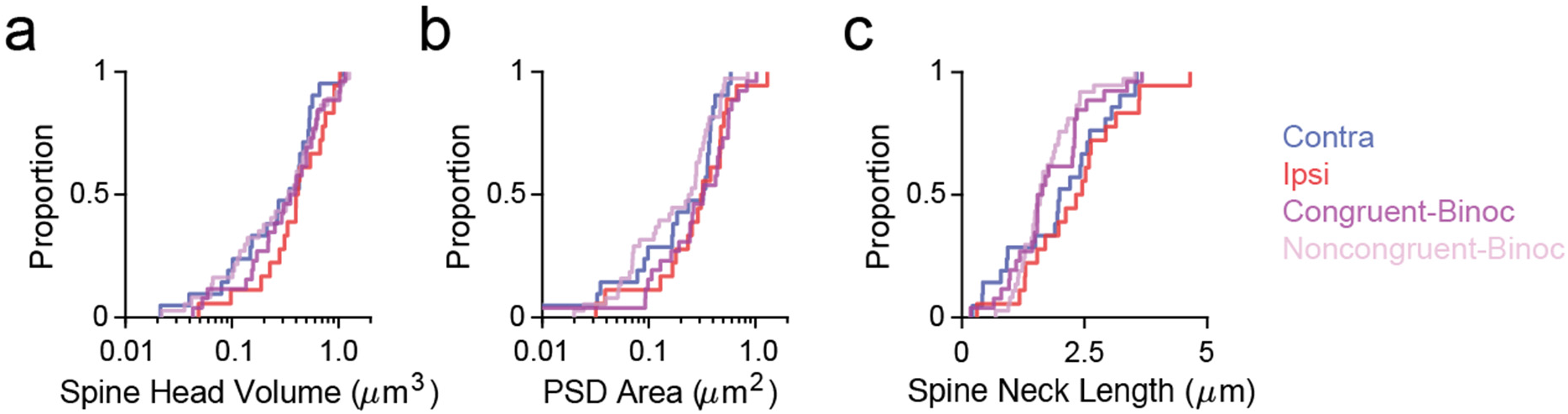
Distributions of synapse anatomy and simulated somatic input for each ocular class. **(a)** Cumulative distributions for spine head volume. Shown are monocular contra spines (blue), monocular ipsi spines (red), binocular congruent spines (purple), and binocular noncongruent spines (light purple). All distributions are statistically indistinguishable. **(b)** Same as in (a) for postsynaptic density area. Binocular congruent spines are significantly larger than noncongruent spines. All other comparisons are not significantly different. **(c)** Same as in (a) for spine neck length. All distributions are statistically indistinguishable.

## Methods

All procedures were performed according to NIH guidelines and approved by the Institutional Animal Care and Use Committee at Max Planck Florida Institute for Neuroscience.

### Viral Injections

Female ferrets (n = 15) aged P18-23 (Marshall Farms) were anesthetized with isoflurane (delivered in O2). Atropine was administered and a 1:1 mixture of lidocaine and bupivacaine was administered SQ. Animals were maintained at an internal temperature of 37 ° Celsius. Under sterile surgical conditions, a small craniotomy (0.8 mm diameter) was made over the visual cortex (7-8 mm lateral and 2-3 mm anterior to lambda). A mixture of diluted AAV1.hSyn.Cre (1:25000 to 1:50000) and AAV1.Syn.FLEX.GCaMP6s (UPenn) was injected (125 - 202.5 nL) through beveled glass micropipettes (10-15 μm outer diameter) at 600, 400, and 200 μm below the pia. Finally, the craniotomy was filled with sterile agarose (Type IIIa, Sigma-Aldrich) and the incision site was sutured.

### Cranial Window

After 3-5 weeks of expression, ferrets were anesthetized with 12.5mg/kg ketamine and isoflurane. Atropine and bupivacaine were administered, animals were placed on a feedback-controlled heating pad to maintain an internal temperature of 37° Celsius, and intubated to be artificially respirated. Isoflurane was delivered throughout the surgical procedure to maintain a surgical plane of anesthesia. An intravenous cannula was placed to deliver fluids. Tidal CO2, external temperature, and internal temperature were continuously monitored. The scalp was retracted and a custom titanium headplate adhered to the skull (Metabond, Parkell). A craniotomy was performed and the dura retracted to reveal the cortex. One piece of custom coverglass (5 mm diameter, 0.7 mm thickness, Warner Instruments) attached to a custom insert using optical adhesive (71, Norland Products) was placed onto the brain to dampen biological motion during imaging. A 1:1 mixture of tropicamide ophthalmic solution (Akorn) and phenylephrine hydrochloride ophthalmic solution (Akorn) was applied to both eyes to dilate the pupils and retract the nictitating membranes. Contact lenses were inserted to protect the eyes. Upon completion of the surgical procedure, isoflurane was gradually reduced and pancuronium (0.2 mg/kg/hr) was delivered IV.

### Visual Stimuli

Visual stimuli were generated using Psychopy (Peirce, 2007). The monitor was placed 25 cm from the animal. Receptive field locations for each cell were hand mapped and the spatial frequency optimized (range: 0.04 to 0.20 cpd). For each soma and dendritic segment, square-wave or sine-wave drifting gratings were presented at 22.5 degree increments to each eye independently (2-3 second duration, 1-2 second ISI, 8-10 trials for each field of view). Drifting gratings of different directions (0 – 337.5°) were presented independently to both eyes. Visual stimulation of either eye was achieved by using a shutter controlled by Arduino which allowed switching between contralateral or ipsilateral view of the monitor at the visuotopic location targeted.

### Two-photon imaging

Two-photon imaging was performed on a Bergamo II microscope (Thorlabs) running Scanimage (Pologruto et al., 2003) (Vidrio Technologies) with 940 nm dispersion-compensated excitation provided by an Insight DS+ (Spectraphysics). For spine imaging, power after the objective was limited to <50 mW. Cells were selected for imaging on the basis of their position relative to large blood vessels, responsiveness to visual stimulation, and lack of prolonged calcium transients resulting from over-expression of GCaMP6s. Images were collected at 30 Hz using bidirectional scanning with 512×512 pixel resolution or with custom ROIs (region of interest; framerate range: 22 - 50 Hz). Somatic imaging was performed with a resolution of 2.05 - 10.24 pixels/ μm. Dendritic spine imaging was performed with a resolution of 6.10 −15.36 pixels/μm.

### Two-Photon Imaging Analysis

Imaging data were excluded from analysis if motion along the z-axis was detected. Dendrite images were corrected for in-plane motion via a 2D cross-correlation based approach in MATLAB or using a piecewise non-rigid motion correction algorithm (Pnevmatikakis and Giovannucci, 2017). ROIs were drawn in ImageJ; dendritic ROIs spanned contiguous dendritic segments and spine ROIs were fit with custom software. Mean pixel values for ROIs were computed over the imaging time series and imported into MATLAB (Sage et al., 2012). Δ*F/F_o_* was computed by computing *F_o_* with time-averaged median or percentile filter (10th percentile). For spine signals, we subtracted a scaled version of the dendritic signal to remove back-propagating action potentials as performed previously (Wilson et al., 2016). For a subset of cells (n = 5/36, n = 1/16 binocular congruent cells), the tuning of the soma was unreliable and an ROI placed on the dendrite apical trunk was used. Δ*F/F_o_* traces were synchronized to stimulus triggers sent from Psychopy and collected by Spike2.

Peak Δ *F/F_o_* responses to bars and gratings were computed using the Fourier analysis to calculate mean and modulation amplitudes for each stimulus presentation, which were summed together. Spines were included for analysis if the mean peak Δ*F/F_o_* for the preferred stimulus was >20% Δ*F/F_o_*, the SNR at the preferred stimulus was > 1, and spines were weakly correlated with the dendritic signal (Spearman’s correlation, r < 0.4). This set of criteria was used to determine the ocular dominance class of each spine and soma. Specifically, if these criteria were passed for stimuli presented only to the contralateral eye (monocular contra), stimuli presented only to the ipsilateral eye (monocular ipsi), or for both (binocular). Binocular congruency was computed as the Pearson’s correlation (Matlab) between mean responses driven by contralateral and ipsilateral stimulation.

Some spine traces contained negative events after subtraction, so response properties were computed ignoring negative Δ*F/F_o_* values. Preferred orientation for each spine was calculated by fitting responses in orientation space with a Gaussian tuning curve (Wilson et al., 2016) using lsqcurvefit (Matlab). Orientation selectivity was computed by calculating the vector strength of mean responses. To identify spine or dendritic calcium events (as used in the model shown in Figures 5-6), ΔF/F traces were smoothed with an exponentially weighted moving average filter (MATLAB) and the peaks of calcium events were located. Peak amplitude of calcium events was compared to the standard deviation of baseline spine fluorescence values prior to subtraction.

### Serial Block-Face Scanning Electron Microscopy

Five layer 2/3 pyramidal neurons from 3 animals previously imaged with *in vivo* two-photon microscopy were imaged with a scanning electron microscope. A total of 23 segments of proximal basal dendrites and 155 spines were reconstructed and analyzed. To facilitate anatomical reconstruction, we limited imaging to proximal basal dendrites. A detailed description of these methods and the data are reported in Scholl et al. (2020) and Thomas et al. (2020).

Fixed (2% paraformaldehyde and 2% glutaraldehyde in a 0.1 mM sodium cacodylate) brain slices of 80 μm thickness were trimmed to approximately 2 × 2 mm to contain the cell of interest at the center. This was accomplished by using blood vessels and slice edges, visible in a 20x epifluorescence image of the slice, as landmarks. The tissue pieces were incubated in an aqueous solution of 2% osmium tetroxide buffered in 0.1 mM sodium cacodylate for 45 minutes at room temperature (RT). Tissue was not rinsed and the osmium solution was replaced with cacodylate buffered 2.5% potassium ferrocyanide for 45 minutes at RT in the dark. Tissue was rinsed with water 2 x 10 minutes, which was repeated between consecutive steps. Tissue was incubated in warm (60°C) aqueous 1% thiocarbohydrizide for 20 minutes, aqueous 1% osmium tetroxide for 45 minutes, and then 1% uranyl acetate in 25% ethanol for 20 minutes. Tissue was rinsed then left in water overnight at 4°C. The following day, tissue was stained with Walton’s lead aspartate for 30 minutes at 60°C. Tissue was then dehydrated in a graded ethanol series (30, 50, 70, 90, 100%), 1:1 ethanol to acetone, then 100% acetone. Tissue was infiltrated using 3:1 acetone to Durcupan resin (Sigma Aldrich) for 2 hours, 1:1 acetone to resin for 2 hours, and 1:3 acetone to resin overnight, then flat embedded in 100% resin on a glass slide and covered with an Aclar sheet at 60°C for 2 days. Since SBF-SEM requires conductive samples to minimize charging during imaging, the tissue was trimmed to less than 1 × 1 mm and one side was exposed using an ultramicrotome (UC7, Leica), then turned downwards to be remounted to a metal pin with conductive silver epoxy (CircuitWorks, CHEMTRONICS).

Tissue was sectioned and imaged using 3View and Digital Micrograph (Gatan Microscopy Suite) installed on a Gemini SEM300 (Carl Zeiss Microscopy LLC.) equipped with an OnPoint BSE detector (Gatan, Inc.). The detector magnification was calibrated within SmartSEM imaging software (Carl Zeiss Microscopy LLC.) and Digital Micrograph with a 500 nm cross line grating standard. Final imaging was performed at 2.0-2.2 kV accelerating voltage, 20 or 30 μm aperture, working distance of ~5 mm, 0.5-1.2 μs pixel dwell time, 5.7-7 nm per pixel, knife speed of 0.1 mm/sec with oscillation, and 56 - 84 nm section thickness. Imaged volumes ranged from 125×125×36 μm to 280×170×52 μm. Serial images were exported as TIFFs to TrakEM2 (Cardona et al., 2012) and aligned using Scale-Invariant Feature Transform image alignment with linear feature correspondences and rigid transformation (Lowe, 2004). Once aligned, each dendrite of interest was cropped from the full volume to reduce computational overhead in subsequent analyses. Aligned images were exported to Microscopy Image Browser (Belevich et al., 2016) for segmentation of dendrites, spines, PSDs, and boutons. Binary labels files were then imported to Amira (versions 6.7, 2019.1) which was used to create 3D surface models of each dendrite, spine, PSD, and bouton. Each reconstructed dendrite was overlaid onto its corresponding two-photon image using Adobe Photoshop for re-identification of individual spines. Amira was used to measure the volume of spine heads and boutons, surface area of PSDs, and spine neck length. Blender (versions 2.79, 2.8) was used to create 3D renderings.

### NEURON Modeling

For each synapse reconstructed, we simulated the change in membrane potential at the soma due to a single action potential arriving at the synapse (on the spine head). Simulations of anatomical features allow generation of a single metric (voltage attenuation between spine head and soma) accounting for a variety of synapse features. We modeled a somatic compartment (radius = 13 μm, *g_Na_* = 0 S/cm^2^, *g_k_* = 0.036 S/cm^2^, *g_leak_* = 0.003 S/cm^2^, *E_leak_* = −50 mV, *R*_a_ = 105 Ωcm, *C_m_* = 1 μF/cm^2^) connected to a 400 μm long dendrite (diameter = 1 μm, *R*_a_ = 105 Ωcm, *C*_m_ = 1 μF/cm^2^). Each spine was placed on the dendrite at the distance from soma as measured with EM and connected via a ‘neck’ to a ‘spine head’ where a synapse was placed. Synapse compartments had the same basic properties (*R_a_* = 250 Ωcm, *Cm* = 1 μF/cm^2^) and passive conductances. Spine neck diameter was fixed to 200 nm, matching widths measured in serial EM sections (data not shown) and the length was set to measured values. Spine head length was set to 1 μm so the diameter could be determined from volume measurements (assigning spine heads to be a cylindrical compartment):

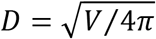

Next, we converted measured PSD area into a value describing the max synaptic conductance. Here we made several assumptions. Based on the linear correlation between the number of receptors and PSD size, we approximated ~0.87 receptors and ~2.0 receptors per 100 nm for AMPA and NMDA, respectively (Takumi et al., 1999). As a simplification, we extract PSD diameter as if our PSDs were circular (as above). Then an AMPA conductance is

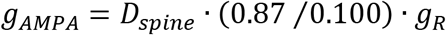

where *g_R_* is 15 pS per channel. In this way, measured PSD area is linearly related to the synaptic conductance used in each model. For each simulation, parameters were set and the maximum depolarization from *V_rest_* (−67.5 mV) was measured in the somatic and spine head compartment.

### Simulation of synaptic population aggregates

To examine the contribution of synapses of different ocular classes we used a simple linear model combining binary visually evoked calcium events. For each binocular congruent cell (n = 14), a random set of synapses were drawn (80% of the total population) and a random set of stimulus trials were drawn for each group of synapses imaged simultaneously on each simulation run (10,000 iterations). The number of active spines was tabulated for each stimulus. Total numbers (or aggregates) were separated by ocular class for comparison. Orientation preference of aggregates was defined as the stimulus evoking the largest response. Multivariate linear regression was used to determine how well total aggregate alignment was predicted by total number of active spines (normalized by the maximum number of active spines observed for each cell) and the fraction of different ocular classes contributing. We used predictor matrices without interaction terms:

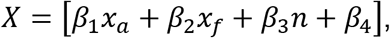

where *x_a_* is the proportion of active synapses at the preferred orientation, *x_f_* is the fraction of spines contributing from a single ocular class, *n* is random noise, and *β* are the linear coefficients. We also used predictor matrices with an interaction term:

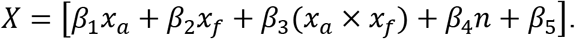

The function fitlm (Matlab) was used to obtain coefficient weights and their significance (see Tables 2 and 3).

### Statistics

Statistical analyses are described in the main text. We used non-parametric statistical analyses (Wilcoxon signrank test, Wilcoxon ranksum test, circular Kruskal-Wallis test) or permutation tests to avoid assumptions about the distributions of the data. All statistical analysis was performed in MATLAB. Correlation coefficients computed with circular-linear correlation or Spearman’s correlation. All correlation significance tests were one-sided unless specified otherwise. Quantitative approaches were not used to determine if the data met the assumptions of the parametric tests.

## Data and code availability

Data and code are available from the corresponding author upon reasonable request.

## Author contributions

B. S. conceived experiments. B.S. and C.Te. performed biological experiments. C.Th. and M.R. preformed electron microscopy and image processing with guidance from N.K.. C. Th., M.R., and B.S. preformed volumetric reconstruction. B.S. and C.Te. analyzed data with guidance from D.F.. B.S. and C.Te. wrote the paper with help from D.F..

